# ImmunoMatch learns and predicts cognate pairing of heavy and light immunoglobulin chains

**DOI:** 10.1101/2025.02.11.637677

**Authors:** Dongjun Guo, Deborah K. Dunn-Walters, Franca Fraternali, Joseph C. F. Ng

## Abstract

The development of stable antibodies formed by compatible heavy (H) and light (L) chain pairs is crucial in both the *in vivo* maturation of antibody-producing cells and the *ex vivo* designs of therapeutic antibodies. We present here a novel machine learning framework, ImmunoMatch, for deciphering the molecular rules governing the pairing of antibody chains. Fine-tuned on an antibody-specific language model, ImmunoMatch learns from paired H and L sequences from single human B cells to distinguish cognate H-L pairs and randomly paired sequences. We find that the predictive performance of ImmunoMatch can be augmented by training separate models on the two types of antibody L chains in humans, *κ* and *λ*, in line with the *in vivo* mechanism of B cell development in the bone marrow. Using ImmunoMatch, we illustrate that refinement of H-L chain pairing is a hallmark of B cell maturation in both healthy and disease conditions. We find further that ImmunoMatch is sensitive to sequence differences at the H-L interface. ImmunoMatch focusses on H-L chain pairing as a specific, under-explored problem in antibody developability, and facilitates the computational assessment and modelling of stably assembled immunoglobulins towards large-scale optimisation of efficacious antibody therapeutics.

## Introduction

The immune system produces an extraordinarily diverse repertoire of antibodies to combat a wide range of immune challenges from invading pathogens as well as endogenous, aberrantly expressed antigens in contexts such as cancer. Our antibody repertoire is diverse, encompassing over 10^12^ distinct antibodies [1, 2, 3, 4]. Such diversity is achieved mainly by two processes: firstly, a random recombination of genetic fragments in the immunoglobulin gene locus occurs independently for the two types of antibody chains, the heavy (H) and light (L) chains [5, 6, 7, 8, 9], which assemble to form an antibody molecule. These recombination processes already generate 2 *×* 10^6^ different combinations of H-L chains [10], the sequence diversity of which is further potentiated by the imprecise joining of these gene fragments [11, 12, 13, 14]. Secondly, mutations accumulated in the antibody variable region exponentiate repertoire diversity [15, 16, 17, 18]. Investigation into the nature, dynamics and regulation of the antibody response *in vivo* [19, 20, 21], as well as the engineering of highly specific antibodies [22, 23, 24], are both active fields of modern biomedical research.

Studies of antibodies have gradually recognised the importance of a wide variety of “developability” factors beyond the binding affinity of the antibody to its antigen [25, 26, 27, 28]. One such issue pertains to thermostability, which is crucial in ensuring the H and L chain can assemble to constitute a manufacturable, functional antibody therapeutic [29, 30, 31]. Using high-throughput sequencing methods, in order to identify stable H-L pairs, researchers generated separate sequencing libraries of H and L chains, manually paired expanded H and L clonotypes, and expressed and tested these antibodies *in vitro* [32, 33]; this has been, for instance, applied to identify broadly neutralising antibodies against HIV-1 [34, 35]. Recent single-cell methods produce paired H-L sequences from thousands of antibody-producing cells [36, 37, 38, 39], which can circumvent problems in the traditional approaches, yet these methods are costly, and still fall short of the true repertoire diversity [40]. The study of how stable antibody H-L pairs are generated is also relevant in basic biology: B cells, precursors to the antibody-secreting plasma cells, express B cell receptors (BCRs) comprising mainly the membrane-bounded version of antibody molecules [41, 42]. During its development in the bone marrow, a B cell undergoes checks to ensure its BCR can be stably assembled and thus can sustain cellular signals to maintain viability, but do not recognise endogenous molecules as antigens [43]. Due to the Darwinian nature of B cell development where cells expressing stable, tolerable BCRs are positively selected [44, 45, 46, 47] and autoreactive B cells are removed via cell death [48, 49, 50, 51]. This poses an inherent challenge in characterising B cells where stable, non-autoreactive H-L pairs fail to be formed *in vivo*. Such knowledge is important to understand the interplay between the formation of a functional BCR and the development of the antibody response, versus the possible autoimmunity arising from defects of this process [52, 53]. Therefore, deciphering the molecular rules governing the pairing of H and L chains will benefit both basic B cell immunology and antibody discovery.

The issue of whether H-L antibody pairing specificity is predictable has been under debate for the past few decades. Seminal antibody structural analyses highlighted the interaction between the H and L chains at the antigen-binding site, comprising mainly of hypervariable regions in each chain known as the complementarity determining region (CDR) (**Figure 1**a) [54, 55]. Contact between the H and L chains is crucial to maintain antibody stability, as well as the orientation of the antigen-binding site [56, 57, 58, 59]. The H-L interface is formed by contacts in the CDR region as well as the antibody framework region (FWR). Molecular biology experiments have identified FWR mutations which would alter H-L interaction geometries and consequently abolish antigen binding [56, 58]. Furthermore, mouse models co-expressing engineered H and L chains often further edit the L chains by introducing mutations which enhance the viability of B cells [60, 61, 62]. Computationally, earlier analyses focused on observations of non-random, over-represented H-L chain partners, however statistical power to identify such associations were limited by the small number of paired H-L sequences and structures available for this type of analysis [63, 64, 65]. The development of single-cell sequencing methods where single B cells in suspension are encased in oil droplets, followed by extraction and sequencing of their H and L chain transcripts [36, 37, 38], has provided new insights to this problem. For instance, statistical analyses in DeKosky et al. [66] suggested that H-L pairing preference was random; however, others argued this could be due to idiosyncrasies in the experimental protocol [38]. More recently, a growing body of evidence suggested coherence between H and L chain choices in B cells. A study by Jaffe et al. [39] using newer single-cell methods found that H chains in mature, antigen-experienced B cells tended to use more restricted L chain partners than their naïve counterparts. Furthermore, comparisons between artifical intelligence (AI) models trained on paired antibody H-L sequences posited that such models trained on paired data outperformed those trained on unpaired chains, in terms of learning biologically interpretable sequence embeddings and predicting antigen specificity [67]; moreover, given the H chain sequence, a stable L chain partner can be generated *de novo* [68]. Taken together, these findings suggest the existence of a set of molecular rules underlying the specificity of antibody H-L pairs. Tools for exploring these rules will hold the promise to understand and design better, more stable antibody pairs, facilitating the development of antibody therapeutics.

**Figure 1:**
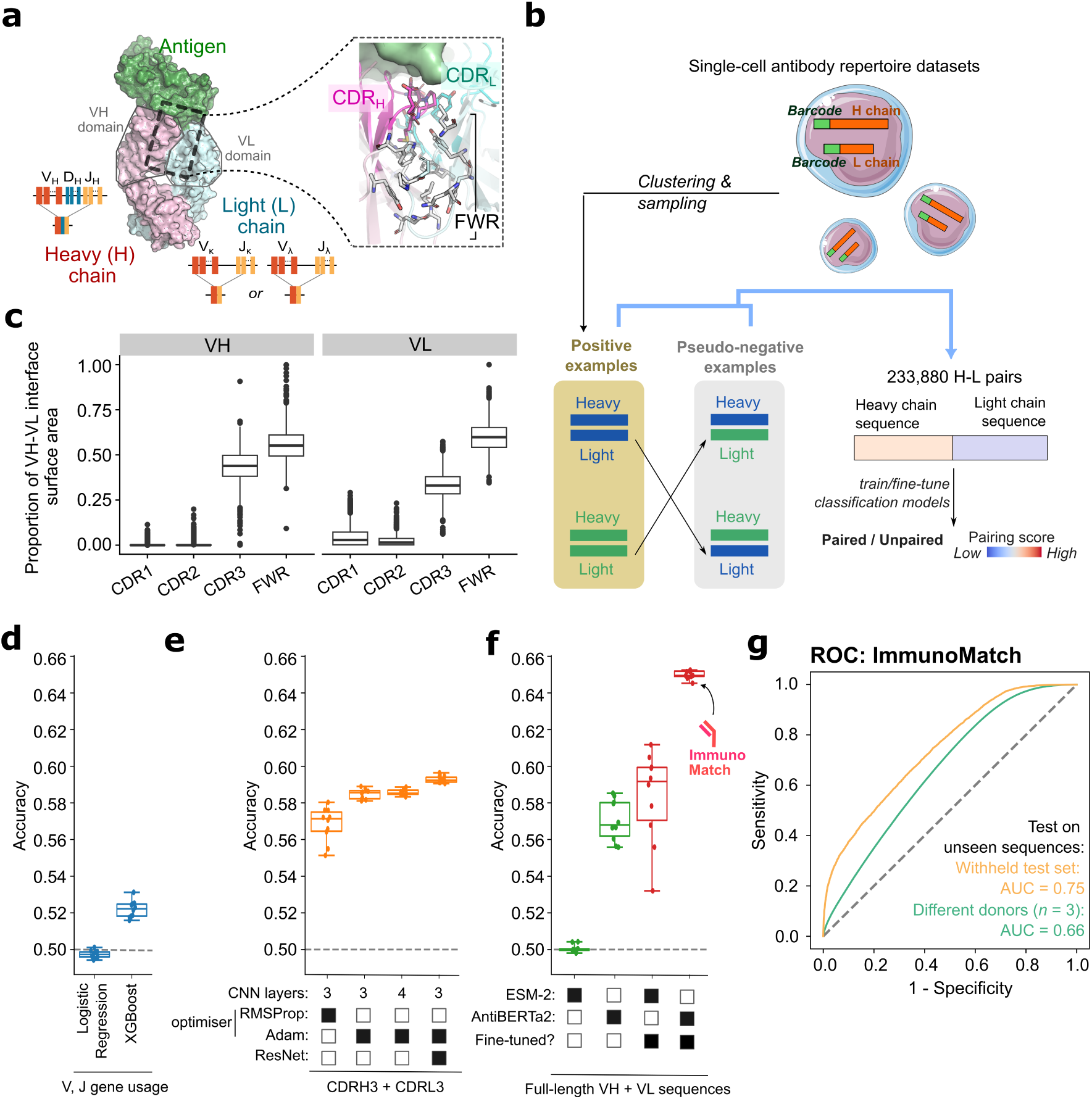
ImmunoMatch for predicting cognate antibody chain pairing. **(a)** Illustration of the antibody heavy (H) and light (L) chain interface (Protein Data Bank [PDB] 6zlr). *In vivo*, H and L chains are generated *via* separate, random recombinations of gene fragments which potentiate the diversity of H-L interface in antibody repertoires. Inset on the right illustrates the H-L interface, with amino acid side-chains (in sticks) to highlight positions in direct contact with the partner chains. CDR, complementarity determining region; FWR, framework region. **(b)** Schematic of model training using curated positive and pseudo-negative examples from single-cell antibody repertoire datasets. See the main text for further details. **(c)** Proportion of the total surface area of the interface formed between the H chain variable region (VH) and the L chain variable region (VL), contributed by individual CDR loops and the FWR, for *n* = 3,781 human antibody structures [70]. **(d-f)** Accuracy of models trained solely on H and L chain variable (V) and joining (J) gene usage (panel **d**), one-hot-encoded CDRH3 and CDRL3 sequences (**e**) and full-length VH and VL sequences (**f**). Each data point represented a separate fold from the 10-fold validation strategy employed during training (see Methods). The final ImmunoMatch model is indicated in panel **f**. **(g)** Receiver operator curves (ROC) of the final ImmunoMatch model, calculated on the withheld test set (area under curve [AUC] = 0.75) and an external test set constituted by *n* = 3 donors unseen during training (AUC = 0.66).

Here, we present ImmunoMatch, a suite of fine-tuned AI models for the classification of cognate antibody chain pairs. Based on an antibody-specific language model (AntiBERTa2 [69]), ImmunoMatch was fine-tuned on full-length H and L chain variable domain sequences extracted from paired antibody repertoire data from healthy donors. This model outperformed baseline models using either CDR sequences or immunoglobulin gene usage as inputs. We found that further optimisation to generate ImmunoMatch variants specific to antibody L chain types improved classification performance. We applied ImmunoMatch to study B cell development through the lens of optimising H-L pairing, and identified chain pairing refinement as a hallmark of B cell maturation in both health and disease. We also validated ImmunoMatch in its ability to recover partner chains in therapeutic antibodies, and highlighted its ability to pinpoint important sequence patterns driving these predictions. Our results underscore the complexity in H-L chain pairing, and highlight the importance of chain pairing in understanding B cell development and engineering stable, functional antibodies.

## Results

### Machine learning models for identifying cognate antibody chain pairing

We posited that machine learning (ML) methods could allow us to test two competing hypotheses, namely that antibody H and L chain pairing preference can be predicted from sequence information, or that such pairing is random. Framing this as a binary classification task to distinguish between cognate, observed H-L pairs from randomly generated pairs, we curated paired H-L sequences, sampled from single-cell antibody repertoire datasets where their cell origin was barcoded as short nucleotide strings [40] (**Figure 1**b). The coexistence of a H and a L chain with the same cell barcode was taken as evidence for paired chains, constituting our positive training examples. Owing to the removal of non-viable H-L pairs by natural selection [43], it was not possible to obtain negative training examples. We instead used a random shuffling strategy to generate “pseudo-negative” examples, exchanging the light chain partners between the observed, positive pairs. This procedure also guaranteed a balanced dataset with equal amount of positive and pseudo-negative examples. Using three separate datasets [66, 38, 39] covering six donors, in total we curated 233,880 H-L pairs for training and testing, after balancing sample sizes over each donor to avoid bias due to the immunological background of individual donors.

We tested the contribution of different input features in combination with multiple ML strategies. Analysing antibody structural data [70, 71], we observed that both the antibody framework region (FWR) and the CDR3 contributed significantly to the interface between the variable heavy (VH) and variable light (VL) domains (**Figure 1**c). Indeed, logistic regression and XGBoost models built solely on V and J gene usage achieved accuracies of 0.50 and 0.52, respectively, indicating limited predictive capability for heavy-light pairing preferences (**Figure 1**d). To improve predictive performance, we next explored using CDR3 sequences as predictive features, in view of its substantial contribution to the VH-VL interface (**Figure 1**c) and their high sequence diversity. We used a one-hot encoding approach and trained a convolutional neural network (CNN) with the CDR3 fragments of the H and L chains, leveraging its ability to capture local patterns within the data. Although the CNN model demonstrated moderate performance, attemps to further improve the model by changing the optimiser, incorporating additional convolutional layers or by adopting the ResNet [72] architecture, yielded minimal improvement (**Figure 1**e). Taken together, these results highlighted the inherent limitations of only considering CDR3 or gene usage, potentially due to the lack of information on specific framework residues that participate in the VH-VL interface (**Figure 1**c).

We therefore explored strategies to incorporate full-length VH and VL sequences in prediction, by capitalising on recent advancement in protein language models [73], as well as those specifically trained using antibody sequences [69]. The transformer architecture, as employed in language models, excels at capturing long-range amino acid interactions [74, 75]. Here, we compared an antibody-specific language model (AntiBERTa2 [69]) against a generic protein language model (ESM-2 [73], 150M parameters) (**Figure 1**f). We observed that by fine-tuning ESM-2, its classification performance increased substantially, comparable to the performance of AntiBERTa2 prior to fine-tuning. The superior performance of AntiBERTa2 suggested that antibody-specific characteristics learned during pre-training were insightful for our task (**Figure 1**f). Through these investigations of different ML architectures (**Figure 1**d-f), we used the fine-tuned model based on AntiBERTa2 as a final instance to classify antibody cognate H-L chain pairing. This model, ImmunoMatch, demonstrated an area under the receiver operator characteristic (AUC-ROC) of 0.75 (**Figure 1**g). To further validate the generalisability of ImmunoMatch, we curated data from *n* = 3 donors unseen during both pre-training and fine-tuning from Jaffe et al [39]. Testing ImmunoMatch on this external evaluation dataset, ImmunoMatch has a AUC-ROC of 0.66 (**Figure 1**g). Details on other performance metrics of ImmunoMatch can be found in **Table 1**. Altogether, this suggests that ImmunoMatch can distinguish cognate antibody VH-VL pairing from randomly paired chains.

**Table 1:** Performance metrics of ImmunoMatch and its variants. MCC, Matthew’s correlation coefficient; AUC-ROC, area under the receiver operating characteristic curve.

| | ImmunoMatch | ImmunoMatch-κ | ImmunoMatch-λ |
| --- | --- | --- | --- |
| F1 score | 0.677 | 0.839 | 0.796 |
| Accuracy | 0.666 | 0.817 | 0.764 |
| MCC | 0.333 | 0.660 | 0.556 |
| Precision | 0.655 | 0.747 | 0.700 |
| Recall | 0.701 | 0.958 | 0.922 |
| AUC-ROC | 0.753 | 0.885 | 0.831 |

### ImmunoMatch performance could be further optimised via a light-chain-specific training strategy

We investigated whether the performance of ImmunoMatch can be further improved. Human antibodies use either one of the two types of light chains, *κ* and *λ*, which are encoded by distinct DNA fragments located on separate chromosomes in the human genome [5]. The VL domains encoded by *κ* and *λ* genes are substantially different, as evidenced by pairwise sequence comparison of *n* = 3,832 antibody structures [70] (**Figure 2**a): on average *κ* VL domains share 47.8% sequence identity with *λ* VL domains, in contrast with the within-group comparisons (69.1% within *κ* light chains, 61.8% within *λ*). We therefore hypothesised that separate models trained on VH-V*κ* and VH-V*λ* sequence pairs could specialise the architecture to learn pairing patterns specific to either type of light chains. Two specialised models, ImmunoMatch-*κ* and ImmunoMatch-*λ* (**Figure 2**b), were fine-tuned using the same workflow as the original ImmunoMatch, with the sole exception that the models were only exposed to light chain of one specific type during training.

**Figure 2:**
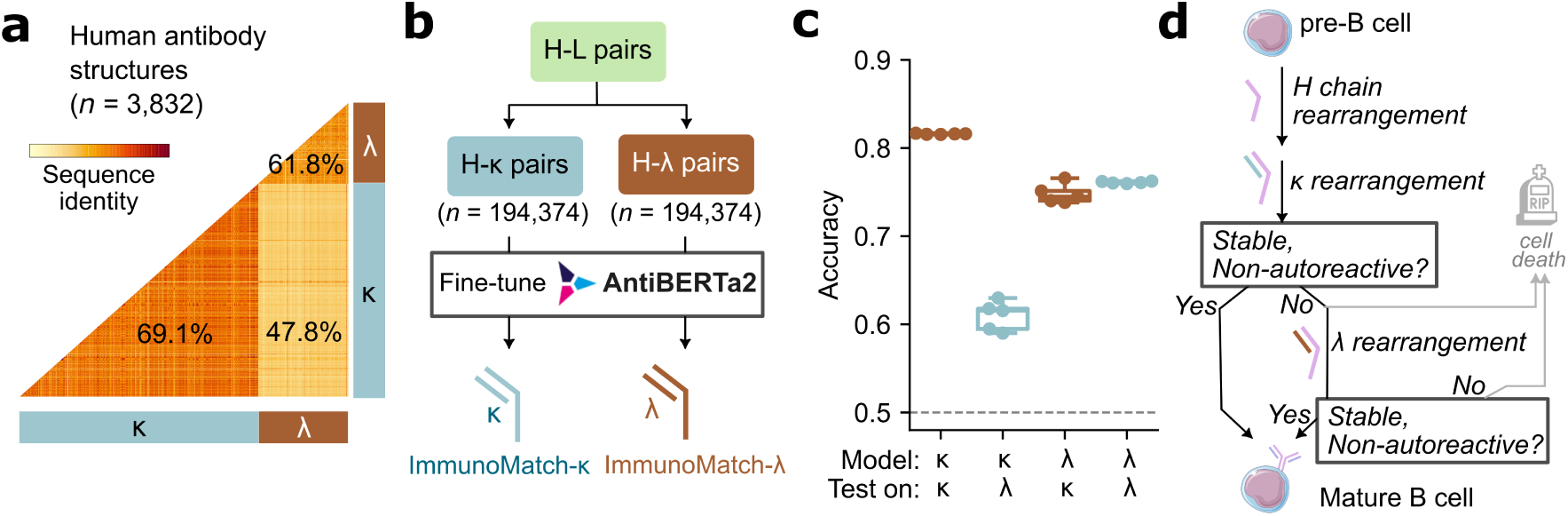
An L chain-specific training strategy of ImmunoMatch was consistent with the *in vivo* mechanism of B cell development. **(a)** Sequence identity comparison between VL sequences taken from human antibody structures (*n* = 3,832) utilising the *κ* and *λ* light chains. The averaged sequence identity within *κ* sequences, within *λ* sequences, and between *κ* and *λ*, are given on the plot. **(b)** Strategy to extract H-*κ* and H-*λ* pairs from publicly available datasets to train separate ImmunoMatch-*κ* and ImmunoMatch-*λ* models. **(c)** Accuracy of ImmunoMatch-*κ* and ImmunoMatch-*λ* on withheld test sets comprised solely of H-*κ* and H-*λ* paired sequences. Each data point corresponded to a separate training fold in a 10-fold cross-validation framework. **(d)** Schematic to illustrate the formation of the B cell receptor *in vivo*. Pre-B cells first rearrange the *κ* locus to generate a stable H-*κ* pair that is not autoreactive, i.e. not recognising self proteins as foreign. Only cells which fail this tolerance checkpoint [43, 76] would either be signalled for cell death, or attempt further *λ* rearrangement.

The performance of these variants of ImmunoMatch was evaluated using separate datasets of antibodies with *κ* and *λ* light chains, which were withheld from fine-tuning. A detailed summary of performance metrics of these models can be found in **Table 1**. ImmunoMatch-*κ* achieved high accuracy (0.817) on *κ* datasets, while ImmunoMatch-*λ* performed comparably well on *λ* datasets, with an accuracy of 0.764 (**Figure 2**c), both substantial improvements over the original Immuno-Match (**Table 1**). When ImmunoMatch-*κ* was tested on *λ* datasets, we observed that this model could still achieve an accuracy above 0.5, albeit with the performance decreased in comparison to the *κ* test set (**Figure 2**c). Interestingly, the performance of ImmunoMatch-*λ* on *κ* datasets remained largely unaffected by the differing distributions of the light chain types between training and testing datasets (**Figure 2**c). This suggests that ImmunoMatch-*λ* is more generalisable in learning pairing rules for antibodies with *κ* and *λ* light chains. This generalisability may be linked to the process of B cell development *in vivo* (**Figure 2**d). Initially, the heavy (H) chains undergo gene rearrangement, followed by the formation of the H-*κ* pairs. They are then subject to central tolerance [43], which either removes B cells expressing unstable and autoreactive pairs of heavy and light chains by signalling them to cell death, or instructs them to rearrange the *λ* gene locus to generate a H-*λ* pair [9, 76, 77, 78]. These H-*λ* pairs are also subject to positive selection of B cells that express a stable H-L chain pair [47, 79, 80] and negative selection of those which react to self-antigens (**Figure 2**d) [76]. Here, compared to the observed H-*λ* pairs, H-*κ* pairs represent a more heterogeneous data source, comprising both H chains which could only specifically pair with *κ* light chains, and those which were ambivalent of the type of light chain partner, but were coupled with a stable *κ* sequence at the first attempt and did not have to undergo *λ* rearrangement. On the other hand, H-*λ* is less heterogeneous, potentially allowing fine-tuning to learn generalisable rules of H-L chain pairing (see Discussion).

### Refinement of immunoglobulin chain pairing is a hallmark of B cell maturation

The analysis above demonstrated that using ImmunoMatch-*κ* and ImmunoMatch-*λ* on H-*κ* and H-*λ* pairs respectively would be more accurate in H-L pairing prediction in comparison to the original ImmunoMatch model (**Table 1**). Using this approach, we next asked whether chain pairing likelihood would vary across stages of B cell development. The classical theory of B cell maturation posits that upon activation, naïve B cells enter the germinal centre (GC) to edit and optimise their B cell receptors (BCR) to specifically bind their cognate antigens, with the successful binders subsequently exiting the GC and differentiating into memory B cells [84, 85, 86]. We collected paired H-L sequences from naïve, GC and memory B cells from published studies [39, 81], and scored the H-*κ* and H-*λ* sequences with ImmunoMatch-*κ* and ImmunoMatch-*λ* respectively. Comparing the pairing scores from these ImmunoMatch models between the B cell subsets, we observed that memory B cells have substantially higher pairing score than their naïve counterparts, with the distribution of GC B cells positioned between the two (**Figure 3**a). We further defined clonally related naïve and memory B cells based on CDRH3 sequence similarity, and identified examples of clonal expansions where pairing score increased as the clonotype diversified from the germline origin (**Figure 3**b). We propose this continuum of chain pairing likelihood to be a feature of B cell maturation: as BCRs undergo class-switch recombination and somatic hypermutation, H-L chain pairing is refined together with these processes, both integral in B cell maturation [17]. To test this hypothesis, we utilised a single-cell RNA sequencing (scRNA-seq) dataset of B cells sampled from the tonsil [81], and compared the H-L pairing score inferred using ImmunoMatch-*κ* and ImmunoMatch-*λ* for sequences of different heavy-chain isotypes and mutational levels. Interestingly, pairing scores increase as the H chain germline identity decreases, and switches away from IgM and IgD isotypes to IgG and IgA (**Figure 3**c). ImmunoMatch pairing scores therefore embed information about B cell maturation, and highlight the increase in H-L pairing specificity as a feature as BCR undergo maturation processes.

**Figure 3:**
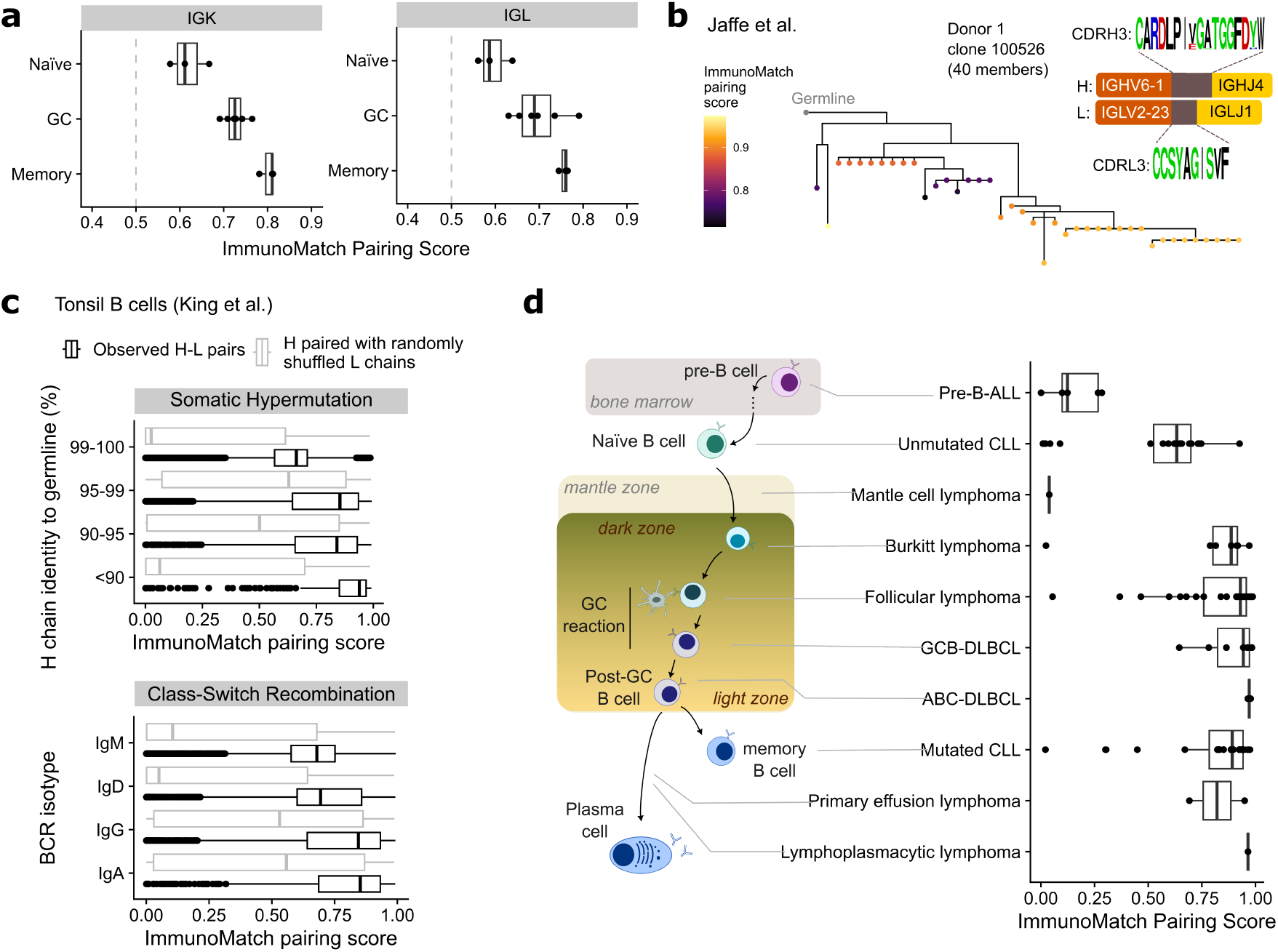
ImmunoMatch revealed a continuum of BCR chain pairing likelihood across B cell development stages. **(a)** Pairing scores of B cell receptors (BCR) from naïve B cells (*n* = 3 donors), germinal centre (GC) B cells (*n* = 6 donors) and memory B cells (*n* = 3 donors). Each data point represents average score per donor, calculated separately for H-*κ* (panel “IGK”, scored using the ImmunoMatch-*κ* model) and H-*λ* (“IGL”, scored using ImmunoMatch-*λ*) pairs. **(b)** Example clonotype tree from Jaffe et al. [39] data with ImmunoMatch pairing scores mapped to individual observations as coloured dots in the tree leaves. The germline configuration of VH and VL sequences are illustrated. **(c)** ImmunoMatch pairing scores for paired VH and VL sequences from tonsil B cells in the King et al. dataset [81], grouped by their germline identity as a proxy of somatic hypermutation status (*top panel*), or by the heavy-chain isotype to illustrate class-switch recombination status (*bottom panel*). As a control, analogous annotation with ImmunoMatch was performed on for each cell, retaining the same observed VH sequence, but each paired with a randomly reshuffled L chain partner. **(d)** Boxplots (right) depicting pairing scores of B cell receptors (BCRs) from leukaemia and lymphoma samples curated from the literature (see Methods). Data were organised by cancer sub-types and ordered by their corresponding B cell development stages when oncogenesis was thought to occur, according to published reviews [82, 83] (schematic on the left). ALL, acute lymphoblastic leukaemia; CLL, chronic lymphocytic leukemia; DLBCL, diffuse large B-cell lymphoma; GCB, germinal centre B-cell; ABC, activated B-cell.

We further investigated whether a similar trend in the pairing scores can be found in BCRs isolated from diseases arising from B cell development. We collected *n* = 123 paired sequences from leukaemia and lymphoma samples collated from the literature and publicly available databases of cancer cell lines, and mapped these samples to the different B cell developmental stages from which these cancers were thought to initiate [82, 83]. Applying ImmunoMatch-*κ* and ImmunoMatch-*λ*, we observed a continuum of H-L pairing scores for these sequences (**Figure 3**d). Specifically, leukaemia originating from pre-B cells in the bone marrow displayed a strikingly low pairing likelihood, reflecting their immature origin [87, 88]. In contrast, in agreement with the need of a functional BCR for B cell activation and antigen interactions in these cancers [89, 90], lymphoma samples typically displayed high pairing scores. These analyses suggest that ImmunoMatch models can be used to annotate immunoglobulin chain pairing, and that the refinement of chain pairing preference is a hallmark of B cell maturation in both health and disease conditions.

### ImmunoMatch is sensitive to sequence differences in therapeutic antibodies

We finally investigated whether our ImmunoMatch mmodels can be applied in an antibody discovery context. Specifically, we simulated an antibody triaging application, where ImmunoMatch-*κ* and ImmunoMatch-*λ* was used to score a random library of germline recombinations of heavy (H)-chain variable (V) and joining (J) gene segments against the cognate light (L)-chain partner (**Figure 4**a). We hypothesised that the best random H-L pair from the ImmunoMatch models should resemble the wild-type sequence. We performed this experiment on *n* = 625 therapeutic antibodies, for which we generated random VH domain sequences, while preserving the observed CDRH3 fragment, for scoring against their cognate VL domains. **Figure 4**b compares the best random VH match against the observed VH in terms of their sequence identities and ImmunoMatch pairing score difference. In line with our hypothesis, the higher the sequence identity, the more likely ImmunoMatch models were to output a similar pairing score. We however noted cases where the pairing score difference was substantial, despite sharing *≥* 80% sequence identity. We identified such cases and mapped the amino acid positions where the randomly generated VH differed from the observed sequence (**Figure 4**c). In these cases, the sequence differences typically reside in the CDRH1 and CDRH2 regions, and also at positions in the framework facing the VL domain (**Figure 4**c). Given the importance of these positions in the VH-VL interface, this highlights the sensitivity of the ImmunoMatch models to structurally important positions in order to infer VH-VL chain pairing.

**Figure 4:**
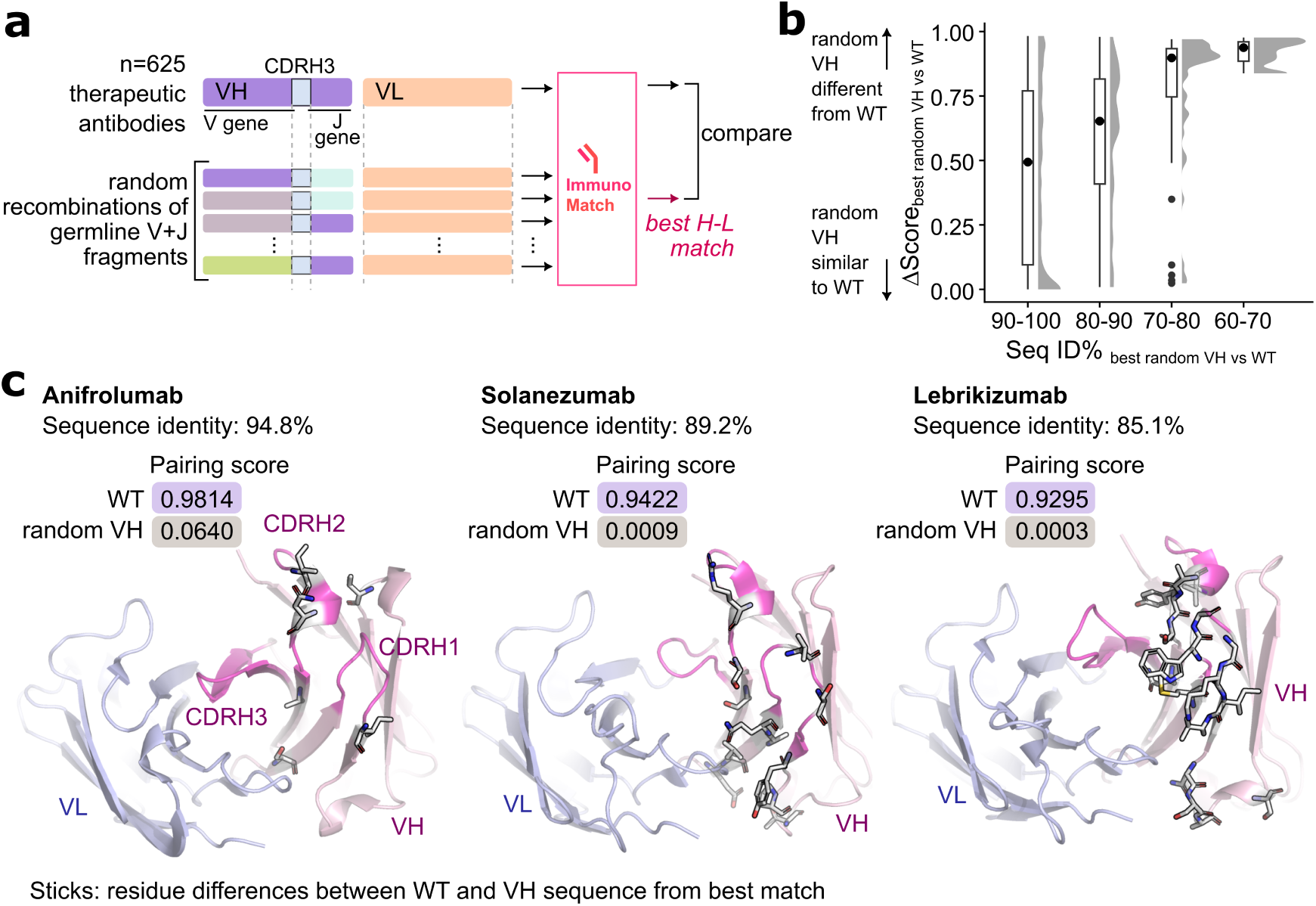
ImmunoMatch is sensitive to VH-VL interface positions in therapeutic antibodies. **(a)** Outline of experiments performed on paired H-L chains from *n* = 625 therapeutic antibodies. For each antibody, we grafted the observed CDRH3 sequence onto random recombinations of H chain germline variable (V) and joining (J) fragments. Together with the observed L chain, these random H chains were subject to scoring by ImmunoMatch-*κ* or Immunomatch-*λ* (depending on L chain type). The random VH sequence from the best H-L match according to the model was compared against the observed VH sequence of the antibody. **(b)** Absolute difference of the pairing scores between the observed (wild-type, or WT) VH for therapeutic antibodies considered in this experiment, and the randomly generated VH ranked best in H-L pairing using ImmunoMatch-*κ* and ImmunoMatch-*λ*. A value of 0 means the pairing scores of the random VH and the WT are identical. The antibodies were binned (horizontal axis) by the sequence identity between the random VH and the WT. **(c)** Examples of therapeutic antibodies where a large pairing score difference was observed between the random H-L pair to the true pair, despite high sequence identity between the VH sequences. Amino acid highlighted in sticks depicted mismatched positions between the random VH and the observed VH used in the therapeutic antibodies.

## Discussion

In this work, we identified an under-explored issue in antibody developability, namely antibody H-L chain pairing, and trained predictive models which specifically address this problem. We found that this was a tractable problem provided the right model architecture was sought, and the resulting modelling framework, ImmunoMatch, demonstrated significant improvement over the baseline and revealed insights into B cell development. ImmunoMatch has several direct, practical uses: firstly, it can be used to assess newly generated single-cell antibody repertoire datasets as quality control, by comparing the ImmunoMatch pairing score distribution of the observed sequence pairs to random background. Given the intensive development of new methods for capturing high-quality paired immunoglobulin sequences from single cells at an increasing throughput [40, 91, 92, 93, 94], a reliable method to assess the validity of the resulting datasets is crucial. Secondly, the ImmunoMatch models can also act as dedicated scoring systems to assess antibody chain pairing, and can be part of a comprehensive assessment of antibody developability. A growing number of computational predictors have been developed to predict antibody solubility, immunogenicity and other characteristics based on their sequences [25, 26, 31, 95, 96]; here, ImmunoMatch can complement the array of existing predictors in order to streamline computational antibody design. Going forward, a unified AI model with a training objective combining these antibody developability measures will be valuable in harmonising the optimisation process of candidate antibody designs. Methods which combine predictions from pools of expert models already exist and have been applied in the evolution of proteins [97]. As antibody development is essentially an engineering process aiming to reach the Pareto front where all desirable characteristics of the antibody are optimised, a combined training objective will facilitate reaching this front by minimising costly iterative processes looking to balance between different developability issues [98, 99]. This is also interesting from a basic B cell biology perspective, since the development of a viable antibody repertoire *in vivo* also follows a similar paradigm, balancing between antigen specificity, cross-reactivity, self tolerance and stably assembled B cell receptors (BCR) capable of sustaining B cell viability [43]. AI approaches which predict these different aspects of BCR biology can be potentially combined to simulate B cell maturation, opening up new avenues in investigating the fundamental pathological mechanisms in diseases where B cell maturation is disrupted.

An important distinction of ImmunoMatch is that its design principles are grounded on basic B cell biology. We reasoned that separate *κ* and *λ* models should produce superior performance compared to the original ImmunoMatch model, supported by the biological process to antibody L chains in the bone marrow. It is surprising that the ImmunoMatch-*λ* model accurately predicted both H-*κ* and H-*λ* pairing; this mirrors the *in vivo* mechanism of B cell development, where the *κ* locus is first rearranged to produce the light chain, followed by *λ* if the H-*κ* pair was eliminated during central tolerance. This secondary, “rescue” role of *λ* light chains during light chain development [100, 101] implies that many H-*λ* pairs are results of failed *κ* rearrangements, potentially facilitating ImmunoMatch-*λ* but not ImmunoMatch-*κ* to learn generalisable rules of chain pairing. Additionally, we designed our randomisation experiments on therapeutic antibodies simulating the assembly of the B cell receptor *in vivo*, screening for successful H-L pairs from libraries of randomly paired antibody chains, and observed a full spectrum of pairing likelihoods given the specifically chosen H and L chains. A number of computational methods aim to simulate antibody repertoires for benchmarking applications, or for learning the statistics of germline gene usage from repertoire data [102, 103, 104]. Our results suggest that simulation without constraining for paired H-L gene usage could significantly oversample dysfunctional sequences incapable of contributing to stable H-L pairs, which would call into question the reliability of these approaches in generating useful benchmarking single-cell, paired-chain datasets resembling real-life antibody repertoires. In general, most approaches to interrogate the functional antibody repertoire have been characterising these molecules at the nucleic acid level, which is known to poorly correlate with protein-level measurements [105, 106]. Going forward, mass spectrometry based antibody repertoire sequencing would allow for direct assessment of observed antibody H-L protein chains [107, 108, 109, 110] to sample well-expressed, stable H-L protein complexes, which is more relevant to our problem.

Predicting H-L chain pairing is highly complex, most notably evidenced by the performance ceiling of the original ImmunoMatch model trained on a mix of H-*κ* and H-*λ* sequences. We have explored possible causes of this limitation and found that this heterogeneity of light chain type is part of the reason, as evidenced by the improvement in performance for the two ImmunoMatch variants specific to either light chain type. However, there are several additional reasons which impose an implicit ceiling to the performance of ImmunoMatch: firstly, due to the limitation in sampling negative examples for training, we devised a strategy to generate pseudo-negative H-L pairs for training. These examples might actually pair *in vivo*, but the incomplete sampling of the repertoire prevented us from observing these pairs. Another possible reason is an inbuilt promiscuity in the number of possible partners of H and L chains. In our positive training examples we observed cases where 1 H chain could have as many as 1,000 distinct L chain partners. Similarly, earlier experimental investigations expressing different combinations of H and L chains often find promiscuous H chains which can be expressed with high yield together with a number of distinct L chains [34, 35, 111]. Whilst we know that antibody germline gene usage is highly biased towards a small handful of genes [112], we do not know whether the stability of H-L chain pairing contributes to skew this distribution. A further extension of our work here would be to investigate the molecular determinants of those examples which are ambivalent towards chain partners, and whether this promiscuity gives such chains a biological advantage to be more represented in the repertoire.

## Acknowledgement

We would like to thank all members of the Fraternali group for comments and suggestions. This work was supported by the Biotechnology and Biological Sciences Research Council [https://bbsrc.ukri.org/, BB/T002212/1 to F.F, D.K.D-W and J.C.F.N.]. D.G. was supported by a PhD scholarship from the China Scholarship Council (number 202008440414). The funders had no role in study design, data collection and analysis, decision to publish, or preparation of the article.

## Methods

### Data curation

#### Data acquisition

Paired heavy and light chain sequences were collected from six datasets across three publications, all derived from healthy human donors (**Table 2**). Data from Rajan et al. [38] and DeKosky et al. [66] were downloaded following their associated publications, and the data from Jaffe et al. [39] were obtained from the Observed Antibody Space (OAS) database [113]. The collected sequences represent diverse B cell populations, including naïve, memory, and mixed B cells. The use of multiple donors served to minimise potential biases in pairing preferences that could arise from donors’ disease states or immunological background. These paired H-L sequences were annotated using IgBLAST [114] to identify germline gene usage and delineate the CDR3 regions for downstream analysis. These collected sequences were subject to the pipeline described below, involving clustering, sampling, and pseudo-negative generation, prior to training and testing machine learning models.

**Table 2:**
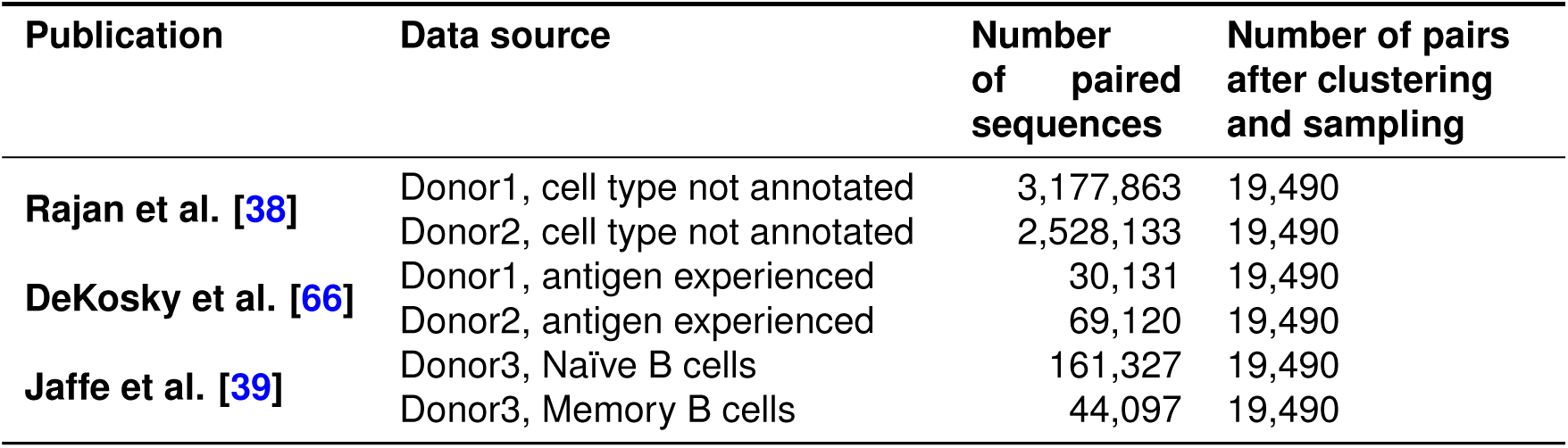
Data collection for paired antibody sequences.

#### Clustering

Clustering was employed to reduce over-representation of specific clonotypes in the training dataset and to prevent data leakage between the training and testing datasets. We extracted paired CDRH3 and CDRL3 (i.e. the most variable antibody segments defining clonotypes [115]), concatenated per pair and clustered them under a 90% sequence identity threshold, using the CD-HIT algorithm [116]. An equal number of heavy and light chain pairs were sampled from each dataset to avoid biases stemming from individual data sources, with details summarized in **Table 2**.

#### Pseudo-negative data simulation

Non-viable heavy and light chain pairs are naturally eliminated during B cell development [43], implying a lack of negative labels to train machine learning models. To address this, we simulated “pseudo-negatives” by randomly selecting two H-L pairs and exchanging their light chain partners. Given the variability in CDRL3 length, this randomisation was further constrained by restricting that only H-L pairs with identical CDRL3 lengths were swapped. This procedure ensured that paired and pseudo-unpaired instances contributed equivalent CDR3 length features, and eliminating signals on H-L chain pairing originating from CDR3 length differences.

Following these steps, the dataset comprises 233,880 sequences with equal amount of positive and negative cases. The pre-processed data are split into training (90%) and testing (10%) subsets for model training and evaluation.

### Baseline models

In establishing all baseline models, these models were trained using a *K*-fold cross-validation approach (*K* = 10).

#### Gene usage-based approach

V and J gene usage was one-hot encoded and used as input for logistic regression and tree-based XGBoost models. Logistic regression was fitted using the LogisticRegressionCV function in scikit-learn (v1.5.0). XGBoost models were fitted using the XG-Boost python package (v2.1.0), with the function XGBClassifier and the parameter “tree_method = ‘hist”’.

#### CDR3-based approach

We leveraged the convolutional neural network (CNN) architecture to capture local patterns in CDRH3 and CDRL3 sequences. Briefly, CDRH3 and CDRL3 amino acid sequences were separately one-hot encoded as a two-dimensional *n × L* matrix, where *n* is the length of the CDR3 fragment and *L* represents the 20 amino acids. The two matrices were passed separately through convolutional layers followed by concatenating outputs from the two streams for passing through to a multilayer perceptron. Various hyperparameters were tested in training the CNN, including the number of convolutional layers, choice of optimizers, and the incorporation of a residual network (ResNet) [72]. For further details, please refer to Supplementary Methods.

#### Language model-based approach

Entire VH and VL paired sequences from the curated training set were passed into pre-trained language models (ESM-2 [150M parameters] [73] and AntiBERTa2 [69]) appended with a classification head for binary classification (paired [positive] *vs.* unpaired [pseudo-negative]). Pre-trained weights of these models were obtained via Hugging-Face. We compared the classification results between the fine-tuned models (for *n* = 3 epochs with learning rate 2 *×* 10*^−^*^5^ and weight decay 0.01, using the Trainer method in HuggingFace) versus the pre-trained models (i.e. weights were frozen during fine-tuning). Please see Supplementary Methods for further details.

### The final ImmunoMatch model

We compared the aforementioned model setups to identify the model with the highest accuracy in the withheld test set. The final model was selected as the fine-tuned AntiBERTa2 model. We call this fine-tuned instance ImmunoMatch. We noted that the withheld test sequences were sampled from the same donors exposed to ImmunoMatch during fine-tuning. We thus curated an external test set, comprising sequences from previously unseen donors in both pre-training and fine-tuning stages. We used paired H-L chains from from donor 1,2 and 4 from Jaffe et al. [39] as this external test set (donor 3 was included in training, see **Table 2**). Data were downloaded from the associated figshare repository (doi: 10.25452/figshare.plus.20338177) of the Jaffe et al. publication. Annotated VH and VL sequences were obtained from the ‘filtered_contig_annotations.csv’ files processed from respective sequencing libraries.

### ImmunoMatch-***κ*** and ImmunoMatch-***λ*** models

We further produced two variants of ImmunoMatch, ImmunoMatch-*κ* and ImmunoMatch-*λ*, by curating antibody heavy-chain sequences with *κ* and *λ* light chains respectively for fine-tuning AntiBERTa2. The previously collected paired antibody sequences were split into two groups according to their light chain types, and separately subject to clustering and pseudo-negatives generation steps. The sizes for training sets of ImmunoMatch-*κ* and ImmunoMatch-*λ* were kept constant (*n* = 194,374). ImmunoMatch-*κ* and ImmunoMatch-*λ* were trained using the same procedure and parameters as the original ImmunoMatch model.

### Applying ImmunoMatch models

For ImmunoMatch, ImmunoMatch-*κ* and ImmunoMatch-*λ*, the epoch with the minimised evaluation loss were loaded for application on external datasets using the HuggingFace “RoFormer-ForSequenceClassification.from_pretrained” interface. A pairing score was derived by applying the softmax transformation on the output obtained by passing through a H-L sequence pair. To apply ImmunoMatch in biological case studies, the annotated sequences were first grouped by their light chain types, and the ImmunoMatch-*κ* and ImmunoMatch-*λ* models were applied on the H-*κ* and H-*λ* subsets respectively.

### Validation datasets

#### Naïve, germinal centre and memory B cell datasets

We collected the following repertoire datasets which comprised paired heavy and light chain sequences traceable to the cell of origin. Firstly, for naïve and memory B cells from healthy individuals, we used data from donors 1, 2 and 4 in Jaffe et al. [39]. Here, only sequencing libraries containing purely sorted naïve or memory (unswitched or switched) were considered. In total we considered n = 711,372 paired sequences from the three donors in the study. Secondly, we curated data from a single-cell study of tonsil B cells [81]. Quality filtered, annotated sequences were downloaded from EMBL-EBI ArrayExpress (accession: E-MTAB-9003) and overlapped with cell-type annotation available for the same dataset based on scRNA-seq [81] (ArrayExpress accession E-MTAB-9005). We used the King et al. dataset for two analyses: first, we extracted sequences corresponding to germinal centre (GC) B cells by considering the associated cell cluster labels (“GC”, “LZ GC”, “DZ GC”, “FCRL2/3high GC”), for comparisons with the naïve and memory B cell data from Jaffe et al. This amounted to *n* = 1,823 paired sequences. Second, we used the entire King et al. dataset in order to examine the differences in ImmunoMatch pairing scores across different heavy-chain isotypes and germline identity levels. For this we considered *n* = 10,782 cells with paired VH and VL sequences, and generated pseudo-negative pairs as control using the identical procedure described in the section “Pseudo-negative data simulation”. Clonotype clustering analysis was performed on the Jaffe et al. dataset using the DefineClones.py function in the Change-O (v1.3.0) [117] package. Clonotype trees were constructed using the maximum parsimony method “dnapars” in PHYLIP [118], accessed through BrepPhylo [119].

#### B cell leukaemia and lymphoma datasets

We compiled paired heavy and light chains from B cell leukaemia and lymphoma samples, from two sources: (1) searches on the GenBank database, which retrieved *n* = 92 paired heavy and light chains obtained from leukaemia and lymphoma samples reported in published works [120, 121, 122, 123, 124, 125, 126], and (2) leukaemia/lymphoma cell lines from the Cancer Cell Line Encyclopaedia (CCLE) project [127]. For (1), sequences were annotated using IgBLAST [114] (v1.19.0) with germline sequences obtained from IMGT (accessed date: 22^nd^ November 2024). For (2), paired-end bulk RNA sequencing FASTQ files were obtained from the Sequence Read Archive (project accession PR-JNA523380), and heavy and light chain sequences were assembled following Tan et al. [128], using the MiXCR (v4.7.0) software [129] run with default parameters. The following filtering steps were performed to ensure paired sequences were from cancers of a B cell origin: (a) removed cell lines with at least 100 reads mapped to T cell receptors; (b) removed cell lines with fewer than 100 reads mapped to immunologubilins; (c) removed cell lines without any heavy chain reads. We noted in these cases, more than one distinct heavy and light chain sequences can often be found [128]. We therefore implemented the following procedure to obtain a single heavy-light chain pair for each cancer cell line: first, we examined the ratio between the total number of reads mapped to the IGK locus to that for the IGL locus; we only considered the specific light chain type if there were at least 100 times more reads mapped to this light chain locus. Second, only the heavy and light chains with the highest fraction of read support were retained. Following removal of out-of-frame sequences we analysed *n* = 31 cancer cell lines from the CCLE project. In total we analysed *n* = 123 paired heavy and light chains from B cell leukaemia and lymphoma.

#### Antibody structure analysis

We analysed *n* = 3,832 human antibody structures curated in VCAb [70]. VH-VL interface surface area was calculated by estimating the change in solvent-exposed surface area (SASA) upon VH-VL complex formation, using POPSCOMP [71]. Definitions of VH and VL domains were taken from VCAb annotations. CDR and FWR regions were delineated using IMGT numbering obtained using ANARCI [130]. For comparisons between *κ* and *λ* light chains, sequence identities were computed using MMseqs2 (v13.45111) [131].

#### Analysis of therapeutic antibodies

We downloaded therapeutic antibody annotations from TheraSAbDab [132] (accessed 5^th^ December 2024), and considered only those which were (1) mono-specific (i.e. with only one unique heavy-light chain pair); (2) approved or under active development; (3) either human or humanised. Sequences were numbered using ANARCI [130]. For each antibody, we kept the observed VL sequence as it was; for VH, we grafted the observed CDRH3 fragment onto all possible combinations of germline *IGHV* - and *IGHJ*-encoded amino acid sequences obtained from IMGT (date: 5^th^ December 2024). This constituted a library of random VH sequences whilst retaining the same CDRH3. This library was screened together with the observed VL using ImmunoMatch to obtain pairing scores, for comparison with the pairing score of the observed VH-VL pair. Sequence identities were computed using MMseqs2 [131] (v13.45111).

## Data availability

Final checkpoints of ImmunoMatch, ImmunoMatch-*κ* and ImmunoMatch-*λ* are available on HuggingFace at https://huggingface.co/fraternalilab/immunomatch. Code to run ImmunoMatch to annotate sequences can be found on Google Collaboratory (https://colab.research.google. com/github/Fraternalilab/ImmunoMatch/blob/main/Run_ImmunoMatch.ipynb). Data and code to generate figures in this manuscript are available on github (https://github.com/Fraternalilab/ ImmunoMatch).

