## Supplementary Methods for "ImmunoMatch learns and predicts cognate pairing of heavy and light immunoglobulin chains"

### S1 Predicting H-L pairing preference using CDR3 sequences

#### S1.1 One-hot encoding of the CDR3 sequences

One-hot encoding is a widely used technique to transform textual data into numerical vectors, enabling subsequent processing by machine learning models [1]. Here, the CDR3 sequences were first one-hot encoded, with each amino acid represented by a vector of length 20, with nineteen `0`s and one `1` at the position of the amino acid in the sequence. For batch processing, sequences were padded with `0`s to achieve uniform lengths. Specifically, CDRH3 sequences were padded to 26 residues, and CDRL3 sequences to 14 residues. The padding strategy adopts the same way as the IMGT-established numbering scheme, where the gaps were inserted in the middle of the CDR3 sequences [2]. After one-hot encoding, sequences were transformed to matrices of shape $(26, 20)$ for the H chain, $(14, 20)$ for L chain and inputted into the CNN model for training.

#### S1.2 Convolutional neural network model architecture

The architecture of the CNN model is illustrated in **Figure S1**. Each pair of one-hot encoded CDRH3 and CDRL3 sequences was individually fed into three convolutional layers, with the first two layers followed by a max pooling layer. Convolutional layers aim to capture the local features of the input data, while max pooling layers aggregate these features by selecting the local maxima and reducing the data dimension. Parameters in convolutional layers were filters being 128, kernel size being 3, with rectified linear unit (ReLU) [3] as the activation function. The setup for the max pooling layer was stride being 3, pooling size being 3. The output of the final convolutional layer was then flattened and processed through four dense layers, with units 256, 256, 128 and 64 and RELu as the activation function. Each dense layer employed a dropout rate of 0.25 to prevent over-fitting. The output was then passed through a dense layer with a single unit and a sigmoid activation function, producing the probability of pairing between the heavy and light chains, resulting in a binary classification indicating whether the chains were paired (using a threshold of 0.5). The CNN model was built using the Keras library in the TensorFlow framework [4].

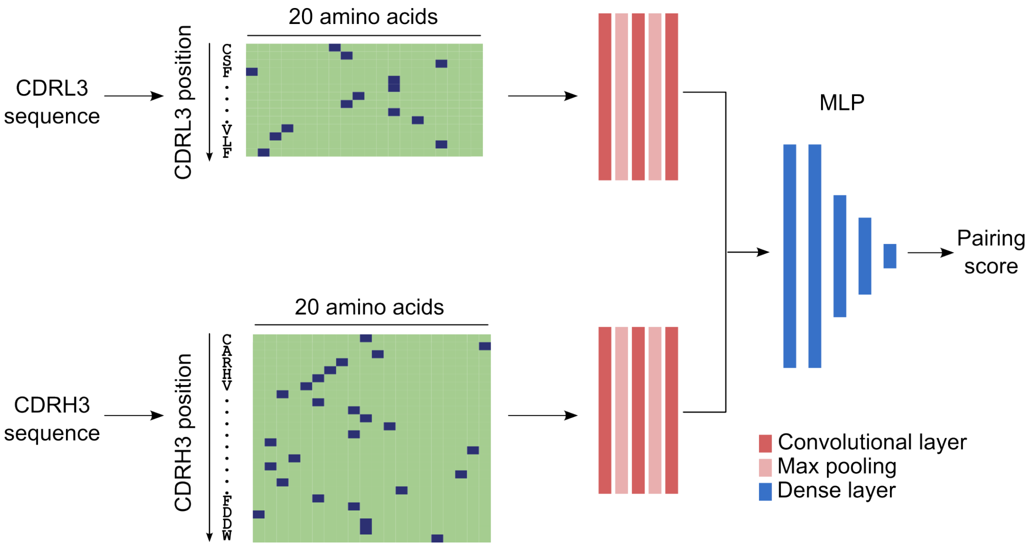


**Figure S1:** The architecture of the CNN model for pairing prediction.
The CDRH3 and CDRL3 sequences were first one-hot encoded as a matrix of shape `(sequence_length, 20)`, with `1`s being the amino acid at the corresponding position (colored blue in the matrix). Encoded sequence were then fed into a series of convolutional and max pooling layers. The output from the CDRH3 and CDRL3 branches were flattened and concatenated, passing through a set of dense layers (Units: 256, 256, 128, and 64; with ReLu as activation function and a dropout rate of 0.25 for each). The last dense layer consisted of a single neuron with a sigmoid activation function, producing the probability of pairing between the heavy and light chains.

### S1.3 Optimisation of CNN model

**Optimiser** The optimiser is a critical component in the training of the neural networks, to update the weights of the model in order to minimise the loss function. Different optimisers have different strategies to update the weights, and the choice of optimiser can have a significant impact on the performance of the model. Here, we investigated the impact of RMSProp [5] and Adam [6] on the performance of the prediction.

**Convolutional layers** Increasing the number of convolutional layers is a common strategy to build a deeper network and improve the performance of the CNN model. Here, the performance of the model using 3 and 4 convolutional layers were evaluated.

**Residual network** Using a deeper neural network by adapting more convolutional layers can improve the performance of the model by allowing the model to learn more complex features. Therefore, we investigated the incorporation of the Residual networks (ResNets) [7] that utilises skip connections between convolutional layers, allowing for a deeper neural network and avoiding the vanishing gradient problem at the same time. Here, the max pooling and the convolutional layer at the end for each CDRH3/CDRL3 branch were replaced by the convolutional and identity block in the ResNet framework (**Figure S2**).

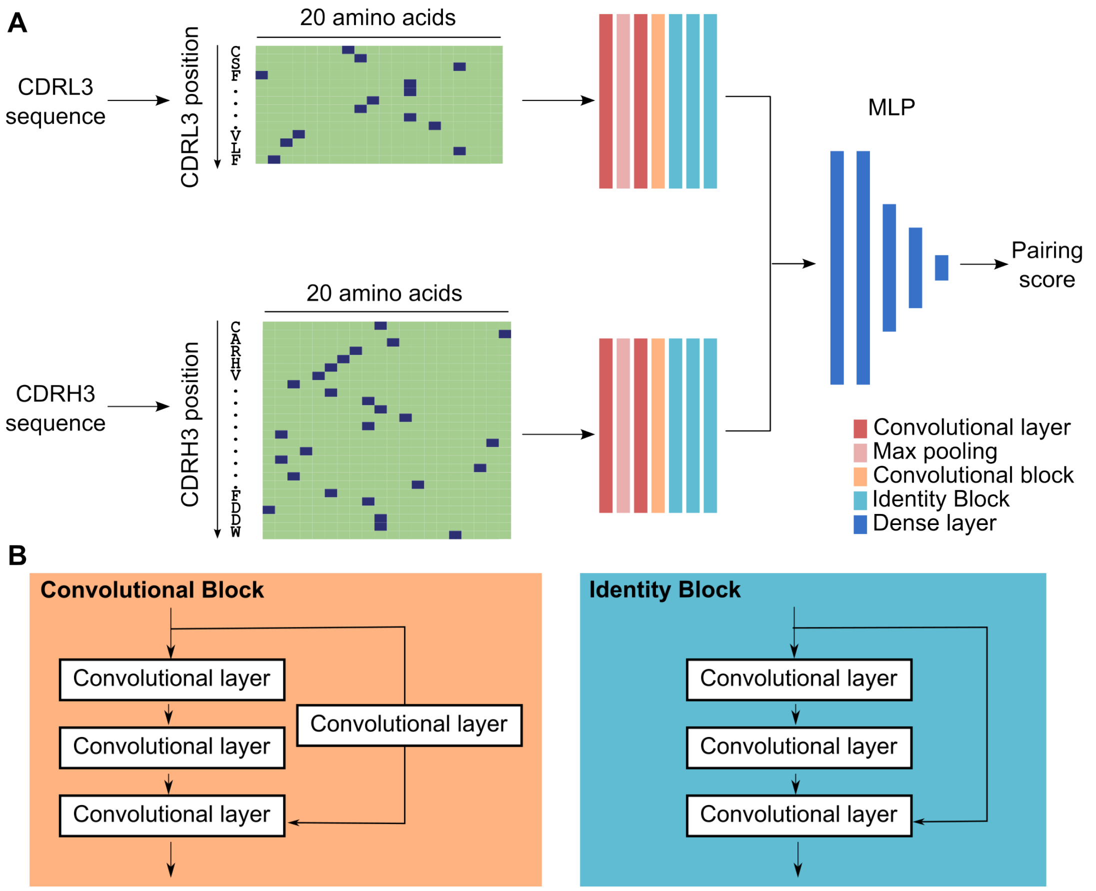


**Figure S2:** CNN model with ResNet architecture.
**(A)** The last max pooling and the convolutional layer in the original CNN structure was replaced with one convolutional block and three identity block. **(B)** The convolutional block consisted of three convolutional layers with a kernel size of 3 and 128 filters, with ReLu activation function. The skip function contained a convolutional layer without activation function, and served to transform the dimensions between the input and output of the block. The identity block consisted of three convolutional layers with a kernel size of 3 and 128 filters with ReLu activation function. A Skip function linked between the input and output of the block.

### S2 Predicting H-L pairing preference using V region sequences

Language models adopt the transformer architecture, where the attention matrix is utilised to better capture the long-range relationships between the tokens in sentences, or, in the case of proteins or specifically antibodies, residues along the amino acid sequence. Since V region sequences ($\sim$ 100 residues) are longer compared with the CDR3 sequences ($max.$ 26 residues), we investigated the use of language models for the prediction of H-L pairing preference using sequences of the entire VH and VL domains as input. Language models can be used as either as a "featuriser", where it encodes the sequence into a fixed-length numeric embedding vector for subsequent machine learning pipelines, or they can also be directly fine-tuned for specific prediction tasks. We investigated both methods in our task, on a generic protein language model (ESM-2 [8]) and an antibody-specific language model (AntiBERTa2 [9]).

### S2.1 Utilising the language model as a featuriser

The VH and VL sequences were separately fed into the language model to output the embedding matrix of shape `(sequence_length, hidden_size)`. Hidden size represents the number of trainable weights for each token, i.e., for each residue. The embedding matrix was then mean-pooled to generate the sentence embedding, where the matrix was averaged along the axis of sequence to produce a fixed-length vector of the shape `(1,hidden_size)`. The two vectors were then concatenated and fed into a multilayer perceptron (MLP) to predict the pairing. The setup for the MLP is the same as in **Figure S1**, with the unit size for the dense layer as 256, 256, 128, 64, and the dropout rate 0.25, respectively. Both ESM-2 [8] and AntiBERTa2 were investigated for this architecture (**Table S1**).

**Table S1:** Language models used in the investigation for pairing prediction.

| Model | Training data | Parameter size | Hidden size |
|---|---|---|---|
| AntiBERTa2 | OAS [10] and antibody structures | 202.64M | 1024 |
| ESM2 | UniRef50 | 150M | 640 |

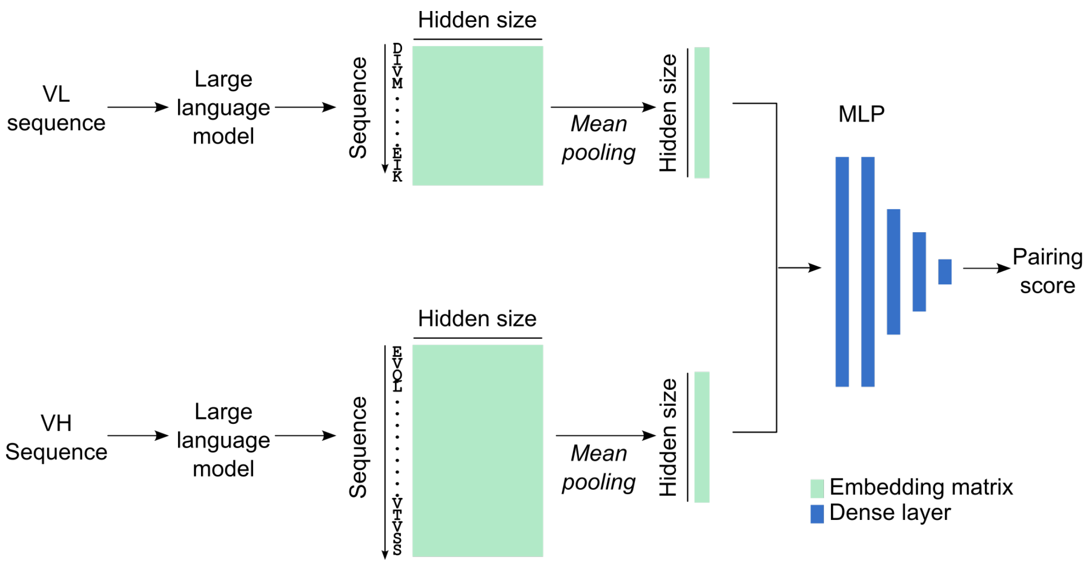


**Figure S3:** Utilising the language model as a featuriser for pairing prediction.
Paired VH and VL sequences were first embedded by different language models, then the embeddings were mean-pooled to generate the sentence embedding. The two vectors were concatenated and fed into MLP (a series of dense layers) to predict the pairing between the heavy and light chains.

#### S2.2 Fine-tuning protein language models for pairing prediction

We also adopted the conventional pipeline to fine-tune ESM-2 and AntiBERTa2 with the training data we collated to predict H-L chain pairing. The VH and VL amino acid sequences were jointly inputted into fine-tuning AntiBERTa2, by adding a separation token (<SEP>) between H and L sequences, a typical method used to concatenate sentences in BERT-based model [11]; for ESM-2, we concatenated VH and VL sequences by adding three gaps (-) in between, as ESM-2 was originally designed for structural prediction [8]. The concatenated sequences were then inputted into the model to fine-tune for three epochs with batch size 48 for AntiBERTa2 and batch size 32 for ESM-2 due to the GPU memory limit and computational efficiency.
